# Orthogonal axes of microbiome variation associated with functionally distinct transcriptomic signatures in the gut of wild *Drosophila melanogaster*

**DOI:** 10.1101/2023.04.24.538093

**Authors:** Frances Llanwarne, Adam J Dobson

## Abstract

Gut microbiota are fundamental for healthy animal function, but the evidence that host function can be predicted from microbiota taxonomy remains equivocal, and natural populations remain understudied compared to laboratory animals. Paired analyses of covariation in microbiota and host parameters are powerful approaches to characterise host-microbiome relationships mechanistically, especially in wild populations of animals that are also lab models, enabling insight into the ecological basis of host function at a molecular and cellular level. The fruitfly *Drosophila melanogaster* is a preeminent model organism, amenable to field investigation by’omic analyses. Here we present an analysis of wild male *D. melanogaster*, with paired measurements of (A) bacterial diversity and abundance, measured by 16S amplicon sequencing; and (B) the host gut transcriptome. We found orthogonal axes of microbial genera, which correspond to differential expression of host genes. The differentially-expressed gene sets were enriched in functions including protein translation, mitochondrial respiration, immunity and reproduction. Each gene set had a distinct functional signature, suggesting that wild flies exhibit a range of distinct axes of functional variation, which correspond to orthogonal axes of microbiome variation. These findings strengthen the bridge between the wild ecology and functional genetics of a leading host-microbiome model.

## Introduction

Animals are ubiquitously associated with microbiota, which play key functions in their hosts’ health (McFall-Ngai et al., 2013). In particular, microbiota in the gut modulate a wide array of host traits, and these functional impacts are observed across the radiation of animals. The penetrance and evolutionary conservation of these impacts suggests that gut bacteria modulate fundamental and ancient host functions, likely via nutrition and immunity (Kawano et al., 2022). This influence means that disruption to the microbial community (aka dysbiosis) can be highly detrimental to host health.

The communities of bacteria that associate with hosts are not fixed among individuals, populations, or taxonomy (Adair et al., 2020). Taxonomic composition of microbiota can vary within and between populations; and also within populations over time (A. C. Wong et al., 2013). However, we do not know if host function covaries with this bacterial variation. Whether or not taxonomically-distinct bacteria can be functionally redundant for hosts remains unknown. Predicting function from taxonomy is challenging even in a single strain of bacteria, because horizontal gene transfer and potentially rapid evolution mean that phylogeny does not necessarily recapitulate function. The situation becomes even more complicated in bacterial consortia, because cooperation and conflict can mean that the emergent properties of a given microbial community may not be predicted by simply summing constituent strains’ functions: for example when interspecific syntrophy enables production of novel metabolites (Consuegra et al., 2020; Henriques et al., 2020); or when one strain’s functions are suppressed by inhibition or competition from other strains (Obadia et al., 2017). In humans, microbiota diversity - defined as taxonomic richness and evenness - has been associated with health and long-term stability of the microbiota (Flores et al., 2014; Huttenhower et al., 2012), but the initial hypothesis that humans harbour a taxonomic core microbiota has given way to a hypothesis of a functional core, in which hosts are thought to need bacteria to serve certain needs, but without reliance on specific taxa (Lloyd-Price et al., 2016). Across the greater diversity of animals, it is similarly unclear whether most hosts associate with a taxonomic core or a functional core. If a taxonomic core is required, then variation may have important functional implications. If a functional core is required, then the impact of variation will depend on the degree of redundancy between fluctuating strains and co-occuring strains.

Appropriate metrics are critical to detect microbial impacts on host function. Metagenomes and metatranscriptomes of microbiota have given valuable insights into microbial genetic competence and gene use (Abubucker et al., 2012; Maurice et al., 2013; Pasolli et al., 2019), but to measure functional impact we must concurrently characterise host traits and processes. The host transcriptome is particularly attractive, because changes to gene expression both underlie and respond to cellular changes (Porcu et al., 2021). The transcriptome is expected to be particularly responsive to microbiota in the intestinal epithelium, since this is the site of physical interaction with gut microbiota. Indeed, the gut transcriptome responds to microbial perturbation in diverse hosts, for example in mice (Aidy et al., 2013; Lavelle et al., 2019; Sommer et al., 2015), *Drosophila* (Bost, Franzenburg, et al., 2018; Bost, Martinson, et al., 2018; Broderick et al., 2014; Erkosar et al., 2014; Fromont et al., 2019), cockroaches (León et al., 2021), and primates (Barr et al., 2018) including humans (Huang et al., 2020). However these studies have tended toward laboratory studies, with consequent limitations of host inbreeding and reduced microbiome diversity. Associating microbiome diversity and function with the host transcriptome stands to circumvent these limitations by examining naturally-occurring bacterial communities, variation therein, and correspondence on function and health of outbred hosts.

The fruitfly *Drosophila melanogaster* (henceforth *Drosophila*) is a well-established system for microbiota studies. In the lab, a major advantage is the simple and culturable *Drosophila* microbiota, dominated by the genera *Acetobacter* and *Levilactobacillus* (Chandler et al., 2011; Staubach et al., 2013; A. C. Wong et al., 2013; C. Wong et al., 2011). Fly embryos can be made germ-free (axenic), and specific bacterial can be reintroduced to make gnotobiotic flies, with a fully-defined microbiota. The same bacterial strains associate with both wild and lab-reared stocks (Chandler et al., 2011), and so ecological studies can potentially be followed by lab investigation (Newell & Douglas, 2014). While *Acetobacter* and *Levilactobacillus* associate with the fly consistently, their abundances fluctuate (A. C. Wong et al., 2013). Other taxa also associate with the fly facultatively, appearing in some surveys but not others (A. C. Wong et al., 2013). These fluctuations have led to a characterisation of the fly microbiota as "inconstant" (A. C. Wong et al., 2013), which may be functionally relevant to the host because distinct strains have distinct functional properties, as shown by both genomic analyses (Newell et al., 2014) and functional assays (Consuegra et al., 2020; Henriques et al., 2020; Leitão-Gonçalves et al., 2017; Newell & Douglas, 2014). Furthermore, the established impact of microbial elimination on the host gut transcriptome (Bost, Franzenburg, et al., 2018; Broderick et al., 2014; Erkosar et al., 2014) leads us to expect transcriptomic correlations to ecological variation in the microbiome. Thus, wild flies and their inconstant microbiota provide an opportunity to investigate whether ecological variation in gut microbiota corresponds to transcriptomic variation in host gut tissues. To date, microbiome-transcriptome associations have been found in congeneric Drosophilids (Bost, Martinson, et al., 2018; Fromont et al., 2019), but not in *D. melanogaster* (Bost, Franzenburg, et al., 2018). Identifying such associations in *D. melanogaster* would potentially enable laboratory studies that leverage the powerful genetic tools available in this system to understand fundamental host-microbe ecology.

Here we present an analysis of variation in the microbiome and transcriptome in the gut of wild-caught male *Drosophila*. The data were published previously, but clear associations were not evident. Here the data are analysed by a distinct approach (Supplementary text), which reveals axes of variation in the microbiota that correspond to host gene expression. Sets of host genes associated with distinct axes of microbial variation were enriched in distinct functions. This analysis suggests that orthogonal axes of ecological variation in microbiome composition and abundance can correspond to equivalently orthogonal axes of variation in host function, in an organism with the capacity to connect natural and laboratory investigation.

## Methods

The study workflow is shown in brief in Figure 1. The full workflow, with details of computational tools, is detailed in Figure S1. Differences between our analysis and the preceding analysis of these same data are presented in the Supplementary Text.

**Figure 1.**
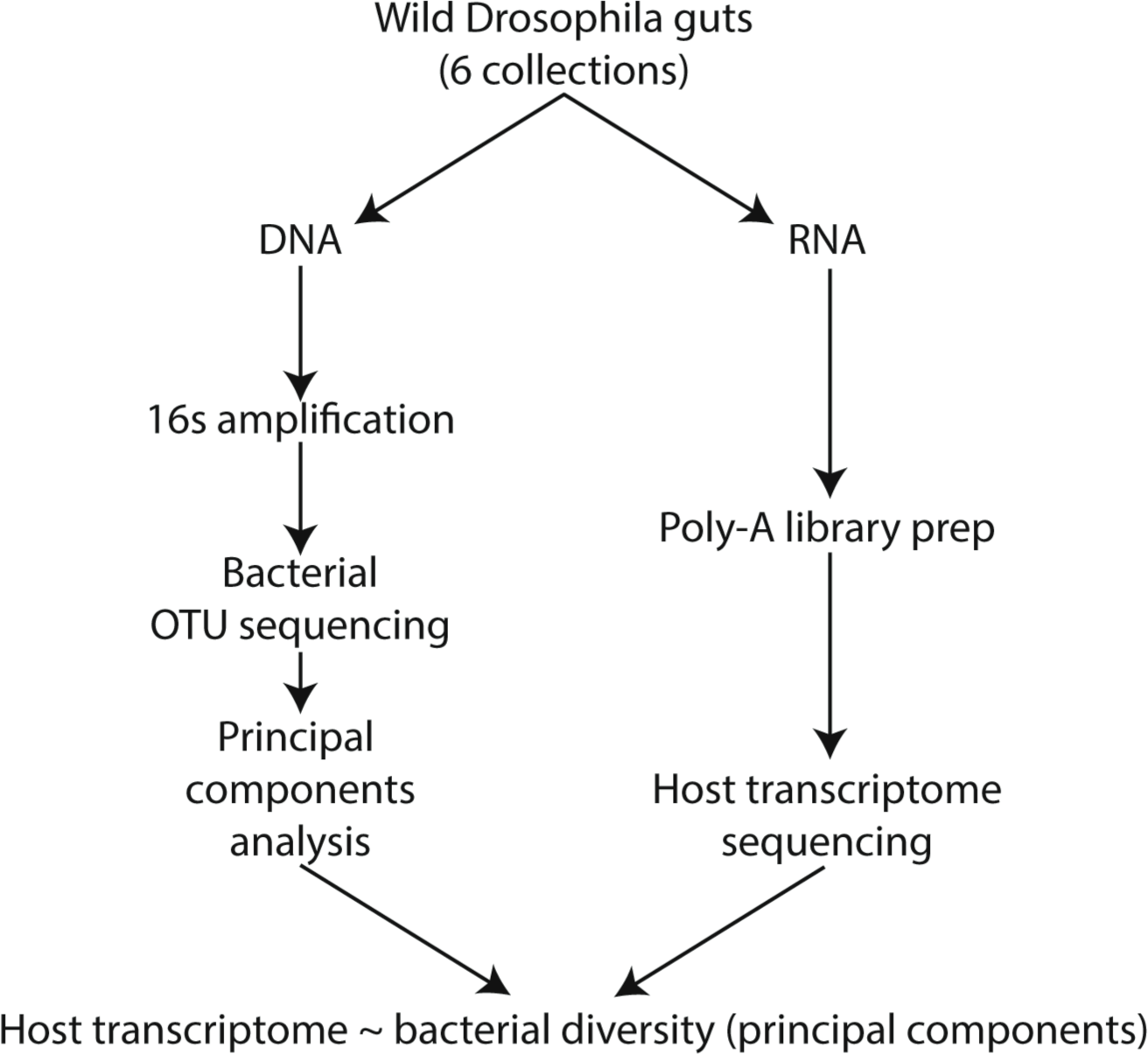
Outline of methodology. Samples were collected by (Bost, Franzenburg, et al., 2018). The resulting public data were reanalysed in this study.

Sample collection methods are described in full by (Bost, Franzenburg, et al., 2018). Briefly, adult wild *D. melanogaster* were collected in Ithaca (samples 1-4) and Geneva (samples 5-6), NY, USA. Females were excluded by necessity, since female *D. melanogaster* could not be differentiated from *D. simulans* by their external features. Dissected guts were subjected to 16S rRNA amplicon sequencing and polyA RNA sequencing, to characterise bacterial diversity and the host transcriptome, respectively. We checked 16S rRNA sequencing quality with FastQC before analysis in QIIME 2 (v 2020.2.0). Sequences were imported into QIIME artifacts (.qza) and the Dada2 plugin was used to conduct quality control with the following adjusted parameters: forward and reverse sequence trimming of the 5’ ends to 6 bp; forward and reverse sequence truncation of the 3’ ends to 260 bp; and pooled chimera checking. Percentages of reads retained after each step are shown in Supplementary Table 1. Closed reference clustering of OTUs was then performed using the vsearch plugin with GreenGenes reference sequences at 94% identity. Chimeric feature sequences were identified and removed (comprising 0.7% of total reads) using the vsearch denovo method. Contaminating endosymbiont OTUs (principally *Wolbachia*) were removed. Taxonomic classification of the resulting features was performed against GreenGenes. First, the desired sequences were extracted from the database using the same primer sequences used for amplifying the bacterial 16S rRNA genes from the sample DNA. A scikit-learn Naïve Bayes classifier was then generated with the extracted reference sequences, and trained with the non-chimeric clustered sequences to create the taxonomy. The taxonomy was then used to collapse the features to level 7 which produced a table with the counts of OTUs per sample. A barplot of the sample taxonomies at a level of either genus, or the next-lowest taxonomic level, was produced in R using ggplot2. Microbial taxonomy was plotted as a Sankey plot in R using the networkD3 library.

*D. melanogaster* transcriptome libraries were downloaded from the NCBI SRA repository using the SRA toolkit. Samples 1-3 and 5 were available as paired-end sequences, while samples 4 and 6 were single-end, therefore, subsequent analyses of the paired-end libraries were performed on the forward reads only. Transcriptome library quality was checked with FastQC. Transcript levels were quantified using Salmon using an index based on reference BDGP6.28 (downloaded from Ensembl). Salmon quantified expression of 14,055 genes, of which 12,911 had at least one read detected in at least one sample. The dataset was filtered to include only genes with ≥1 read in each sample, leaving 8,960 genes for input into further analyses. Library depth and numbers of genes before and after filtering are reported in Supplementary Table 1.

Subsequent quantitative analyses were performed in R (v4.2.1). OTUs with fewer than 4 reads among all samples were excluded, and OTU counts were transformed by natural logarithm. PCA and factor mining were performed using FactoMineR::PCA and FactoMineR::dimDesc (p≤0.05). Genes with zero read counts across all samples were excluded. Coefficients of variation were calculated on rLog-transformed reads. Heatmaps of gene expression counts (rLog-transformed, scaled and centred) were plotted with the Superheat package. Differential gene expression was determined using DESeq2. OTU PCs 1-4 were mean-centred and scaled, then included as predictive variables in a DESeq2 model fit. Impact of each PC was determined by a likelihood ratio test when each respective PC was excluded from the model. Genes were accepted as differentially expressed when at a false discovery rate ≤0.01. Gene Ontology (GO) enrichment was analysed using TopGO, using the "classic" algorithm to run Fisher’s test. GO dotplots were plotted in ggplot2.

Figures were assembled in Adobe Illustrator v26.5.

## Results

### Microbiome diversity

We first examined microbial diversity in the wild fly gut samples. We found a total of 99 operational taxonomic units (OTUs, Figure 2a). In accordance with previous surveys of wild flies (Chandler et al., 2011; Staubach et al., 2013; C. Wong et al., 2011), the family *Acetobacteraceae*, which are key for fly function (Henriques et al., 2020; Leitão-Gonçalves et al., 2017; Newell & Douglas, 2014), were most abundant (Figure 2a). Of the six samples, five were indeed dominated by *Acetobacteraceae*, comprising up to 95% of per-sample OTU counts (Figure 2b). The remaining sample (sample 3) was dominated by *Enterobacteriaceae*, comprising 94% of OTU counts (Figure 2b). Per-sample numbers of distinct OTUs, counts, and alpha diversity, were all variable (Figure 2c): numbers of distinct OTUs varied from 20 (sample 2) to 64 (sample 6); OTU counts ranged from 8,418 (sample 5), to 33,062 (sample 4); and alpha diversity (Shannon index) ranged from 0.313 (sample 4) to 1.886 (sample 5).

**Figure 2.**
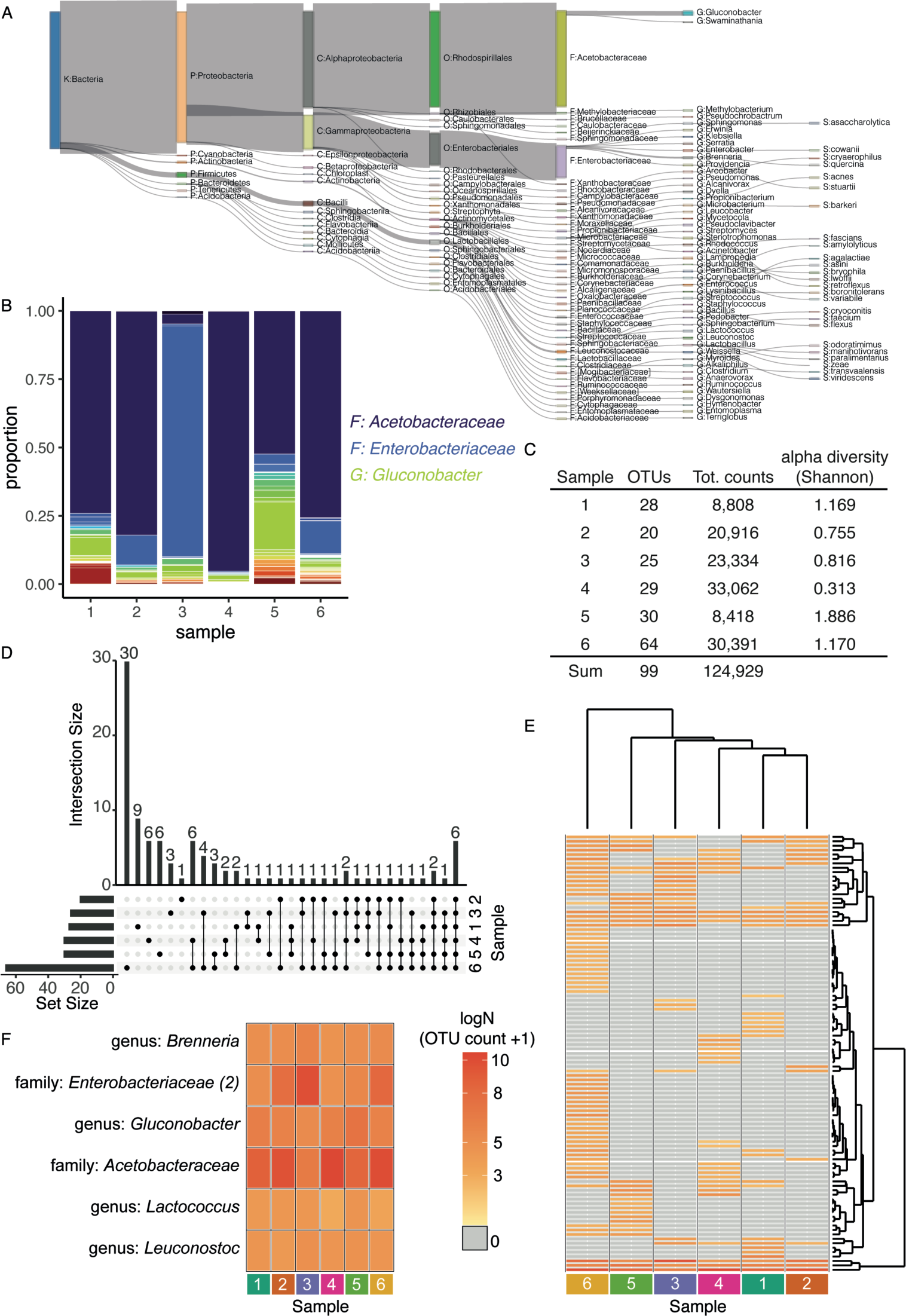
Microbiome diversity in wild male *Drosophila* guts. **(A)** The taxonomy of bacteria identified by 16S rRNA sequencing and analysis with QIIME (dada2 & GreenGenes). Sankey plot shows hierarchy of OTUs detected in the wild fly gut samples. For each level of taxonomy, widths of edges are scaled to proportion of reads assigned to the given OTU, e.g. 100% of reads were from the kingdom Bacteria; the majority were then subclassified as phylum Proteobacteria; which were then mostly sub-classified as class Alphaproteobacteria then Gammaproteobacteria, etc. Branches terminate at the lowest taxonomic level that could be confidently called. **(B)** Relative microbiome composition among samples. Barplot shows proportional representation of OTUs in each per sample. Key shows colour-coding of the three most abundant taxa. **(C)** Variation in within-sample index of diversity. Table shows Alpha diversity (Shannon index) per sample. **(D)** Varying degrees of intersection among samples in OTUs. Upset plot shows intersections of OTUs per sample. Left-side barplot shows total OTU counts per sample. Sample ID given to right of matrix. Connected dots in matrix designate intersections between samples. Top barplot shows size of the intersection. **(E)** Flies varied by OTU occurrence and abundance. Heatmap showing occurrence and abundance of OTUs per sample. Samples and rows ordered by hierarchical clustering, shown by dendrograms above and to right. Colour key shown to left (shared with panel F). Absence of OTU (i.e. 0) denoted by grey colouring. **(F)** OTUs identified in all samples vary in abundance. OTU identity is given to left, colour key shown to right (shared with panel E).

*Acetobacteraceae* are a leading driver of fly function. While *Acetobacteraceae* OTUs were abundant, QIIME could not distinguish strains for most counts (Figure 2a). However, previous genomic analysis of fly-associated strains suggested that most *Acetobacter* strains are commonly competent for similar biosynthetic pathways (Newell et al., 2014), and so we continued with our analysis, binning most *Acetobacteraceae* into one family-level OTU.

To evaluate among-sample diversity, occurrence and abundance of OTUs per sample were compared. Intersections between sets of OTUs per sample were plotted (Figure 2d), revealing six OTUs that were present in all six samples. Each sample also contained unique OTUs, the largest set of which comprised 30 OTUs uniquely assigned to sample 6 (Figure 2d), whereas sample 2 contained only one unique OTU (Figure 2d). 39 OTUs occurred in intermediately-sized sets among 2, 3, 4 or 5 of the samples (Figure 2d).

To visualise the distribution of OTU occurrence and abundance, counts per sample were hierarchically clustered and plotted in a heatmap (Figure 2e). Sample 6 clustered away from other samples, in accordance with its high frequency of unique OTUs. Samples 1-5 clustered more closely. To study abundance of the putative core microbiota specifically, a second heatmap was plotted comprising just the six core OTUs (Figure 2f), revealing variable among-sample abundance.

Altogether, the 16S rRNA analysis indicated among-sample variation in the microbiome, with ecological variation in microbiome taxonomy.

### Transcriptome diversity

Having established among-sample variation in the microbiome, we asked whether the samples also exhibited variation in the transcriptome. A total of 74M transcripts were quantified, and mapped to 8,960 genes. To assess quantitatively the among-sample variation in transcription, coefficients of variation (i.e. variance/mean) were calculated for expression of each gene, and expression of the top 1000-ranking genes were plotted in a heatmap with hierarchical clustering (Figure 3). The heatmap indicated clusters of genes with distinct expression levels among the six samples. Thus, the samples under study exhibited ecological variation in both the microbiome and transcriptome.

**Figure 3.**
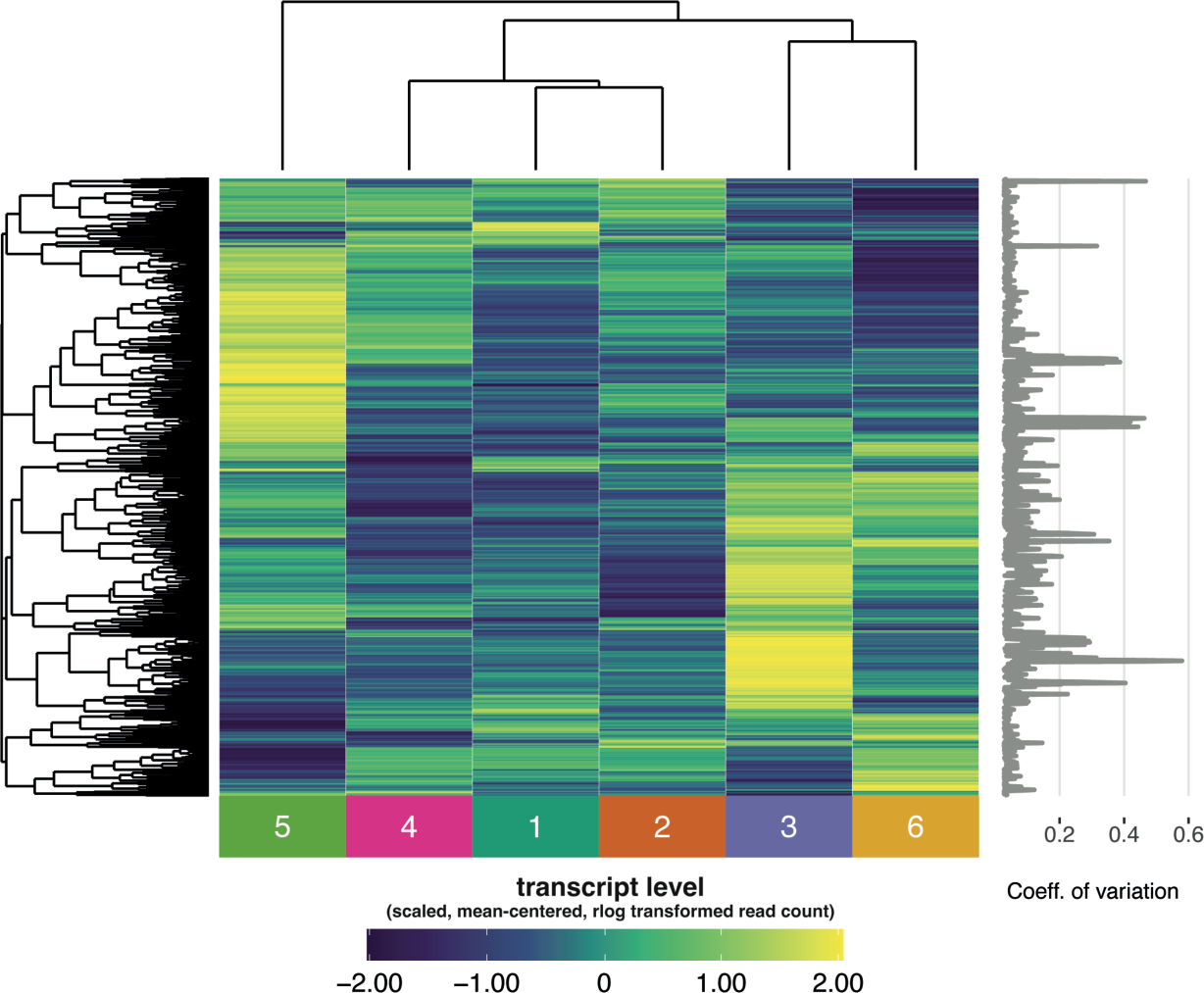
Transcriptome diversity in wild male *Drosophila* guts. Heatmap analysis suggests transcriptomic variation in the wild fly gut samples. The heatmap shows per-sample read counts (rlog scale) for genes with the 1000 highest-ranking coefficients of variation (shown to right). Read counts are scaled and mean-centred to row. Sample identities are given at the bottom. Rows and columns are ordered by hierarchical clustering, as shown by dendrograms.

### Principal components of microbiome variation

We sought an aggregate view of the major axes of variation in the microbiota, to provide predictive variables to parse host transcriptome variation. We were interested to ask whether microbiota taxonomy predicted host gene expression. We reasoned that mapping the host transcriptome to the lowest available level of bacterial taxonomy in our 16s analysis (species) would not be informative, as the distribution of OTUs would be too granular, and indeed previous genomic analysis showed close similarities in the genomic competence of congeneric bacteria (Newell et al., 2014). We therefore characterised axes of microbial variation at the level of genus, or the next-lowest taxonomic level that a given OTU could be mapped to. The OTU table was filtered for OTUs with at least four counts at the lowest available level among the six samples (which excluded 5 OTUs). Counts were then summed per OTU at the level of genus or above, log-transformed, scaled, and submitted to principal components analysis (PCA). A biplot of the first two PCs indicated evenly-distributed loading of OTUs (Figure S2a). The first PC explained 43% of total OTU variance, while PCs 2-4 each explained between 14-18% (Figure S2b). We excluded PC5 from further analysis, because of the relatively low proportion of variance explained (9%, Figure S2B). Pairwise plots confirmed that the PC values were orthogonal (Figure S2C). We also noted that PC1 strongly differentiated sample 6 from other samples (Figure S2C), consistent with its distinct OTU profile (Figure 2c-e). To assess whether the PCs represented real axes of microbiome diversity, we reverse-associated PC values with OTU counts from the original dataset using factor mining tools, plotting values of each PC and presence/absence and abundance of its associated OTUs in a heatmap, ordered by PC values (Figure S3). Each PC had corresponding OTUs, in sets of 32 to 9 per PC (Figure S3). This confirmed that the microbiota PCA revealed orthogonal axes of aggregate variation in the microbiota, amenable to tests for association with the associated host gut transcriptomes.

### Transcriptome variation associated with principal components of microbiome variation

Are axes of variation in the microbiome and the transcriptome correlated? We used DESeq2 to test for correspondence of host gene expression to microbiome PCs. For each PC, a corresponding set of differentially expressed host genes was identified. Sets of 33, 33, 51, and 6 host genes were associated to PCs 1-4, respectively (Supplementary Tables 2-5). To visualise this covariation, we plotted gene expression and associated PC values in heatmaps, ordered by PC values. These plots confirmed quantitative covariation between microbiome PCs and the host transcriptome (Figure 4). We note that the divergence of the microbiome in sample 6 drove a highly divergent value for this sample on PC1, and so these results must be treated conservatively. Altogether, these data indicate that ecological variation in the taxonomic composition of the wild fly microbiota can correspond to host gut gene expression.

**Figure 4.**
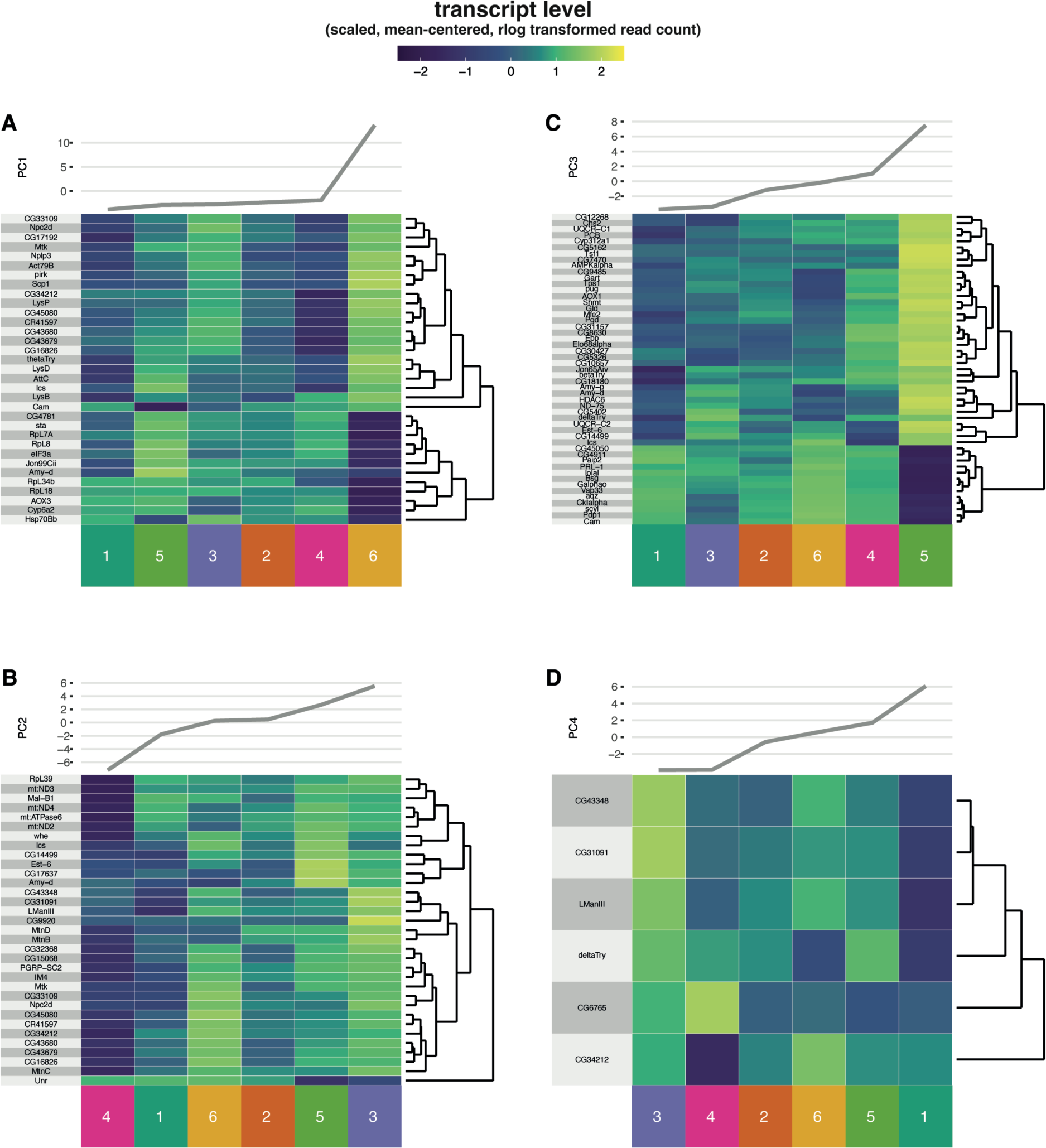
Host gene expression corresponds to axes of OTU variation in the microbiota. Major axes of variation in OTU counts were identified by principal components analysis (PCA, Supplementary Figures 2 & 3). Resulting PCs were used as predictive variables in differential expression analysis of host transcriptome, identifying sets of genes associated with each microbiota PC. Panels **A-D** show heatmap analysis for genes associated to microbiome PCs 1-4, respectively. Heatmaps show expression values (rlog scale) for associated gene sets. Line plots to top of each panel show values of each respective microbiome PC. Read counts are scaled and mean-centred to row. Sample identities are given at the bottom. Samples are ordered by microbiota PC values (at top). Rows are ordered by hierarchical clustering (dendrograms to right of figures). The correspondence between heatmap colouring and PC lineplots indicates patterns of gene expression that correlate patterns of ecological variation in the microbiota.

### Functional analysis of transcripts associated with microbiome variation

Which host functions correlate ecological variation in the microbiota? We analysed enrichment of Gene Ontology (GO) terms amongst differentially-expressed transcripts, for the union of sets of differentially-expressed genes associated with microbiota PCs 1-4 (Supplementary Table 6). We analysed ontologies for biological process (Figure S4), molecular function (Figure S5) and cellular component (Figure S6) separately. The union of gene sets was enriched in biological functions related broadly to immunity, reproduction, metabolism and mitochondrial respiration, protein translation, and immunity (Figure S5, Supplementary Table 6).

Could specific sets of taxa in the microbiome modulate specific host functions? If so, then taxonomically orthogonal axes of microbiome variation might correspond to expression of sets of host genes with distinct functions. We therefore assessed enrichment of GO terms for each respective set of host genes associated to microbiome PCs 1-4 (Supplementary Tables 7-10). Per GO term, we plotted p-values, total number of annotated genes, and proportion of observed/expected genes, for the top 5 GO terms per gene set (Figure 5), which suggested that each set of host genes was associated with discrete sets of GO terms (Figure 5). Expanding this plot to the full set of genes gave the same signal of distinct functions associated with each distinct gene set (Figure S7). Thus, these results suggest that taxonomic variation in the microbiota correlates differential regulation of specific host gut functions, generating complex intersections of functional diversity.

**Figure 5.**
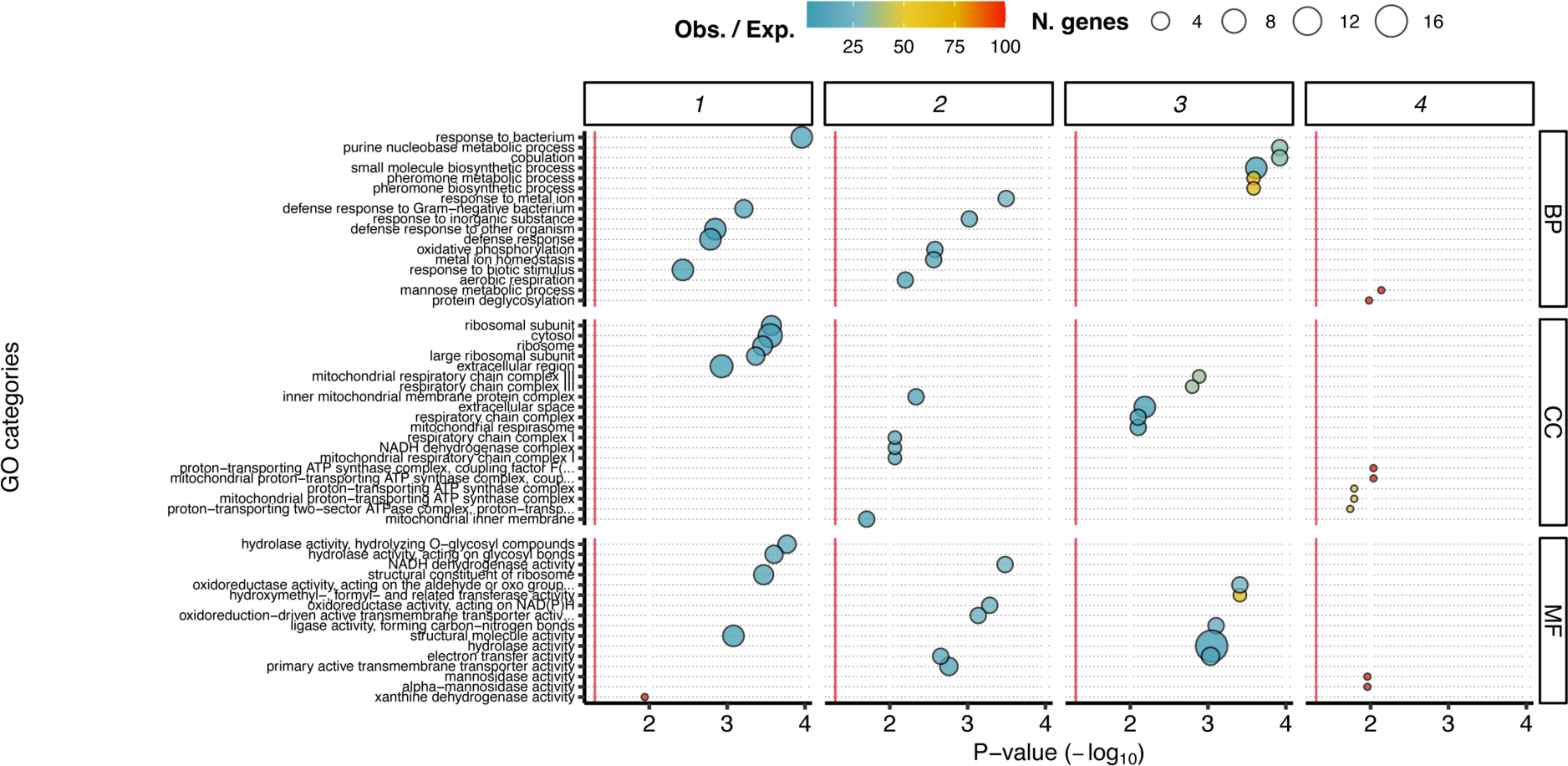
Distinct host functions associated to orthogonal axes of microbiota variation. Gene ontology enrichment was calculated for gene sets associated to microbiome PCs 1-4, for biological process (BP), cellular component (CC) and molecular function (MF) ontologies. Panel shows enrichment per gene set (columns) and ontology (rows). Bubble plots show enrichment of GO terms per PC and per ontology. X axis shows P-values (Fisher’s test, −log_10_ scale) for enrichment. GO terms are given on Y axis, ordered by average of X axis values across all gene sets. Bubble size shows number of genes in the set associated with the GO term, colour shows ratio of n. observed/expected genes. The top-5 ranked GO terms per each gene set are shown. See Supplementary Tables 7-10 for detail of GO terms, and Figure S7 for an equivalent plot of the full set of GO terms.

## Discussion

This study shows that expression of the host gut transcriptome can correspond to microbiota taxonomy in a wild animal. Specifically we have shown that variation in microbiota OTUs at the level of genus or higher correlates to host gene expression. Orthogonal axes of microbial diversity are correlated to sets of genes with distinct host functions. Those functions altogether are consistent with established effects of the gut microbiota on gut function.

Gut microbiota are well-established regulators of the host transcriptome. However the majority of studies have been conducted on laboratory systems, in which microbiome diversity is typically diminished, and hosts are typically inbred (Fromont et al., 2019). Furthermore, laboratory systems have tended to quantify differences in dyadic analyses of animals reared with or without microbiota, providing no scope to correlate differences in microbiome abundance and diversity with transcriptomic differences. Analyses of the correspondence between the microbiome and transcriptome in wild animals, such as that presented in this manuscript (Bost, Franzenburg, et al., 2018; Bost, Martinson, et al., 2018; Fromont et al., 2019), provide the opportunity to address this gap. Our finding of an association suggests that equivalent patterns may be discernable in hosts with more complex microbiota. This task is relatively trivial if the microbiota exist in discrete states, e.g. "enterotypes" (Arumugam et al., 2011), largely because it is straightforward to identify differential gene expression between discrete groups. However there is an argument that the concept of enterotypes over-simplifies the microbiome, and that gut microbiota vary along continuous axes (Knights et al., 2014). Our analysis shows that impacts of continuous microbial variation on gene expression can be modelled by using outputs of dimension-reduction techniques, like PCA, as inputs into differential expression analysis.

Our analyses indicate a diverse array of host processes that correlate variation in the microbiota. This suggests a degree of genetic modularity, with the bacteria generating quantitative variation in host function. Some of the functions indicated are already known to respond to microbiota, for example immunity (Buchon et al., 2009) and mitochondrial respiration (Gnainsky et al., 2021). Other implicated functions are less well associated to the microbiota, for example translation. The suggestion that these functions vary in wild flies in concert with microbiota suggests that they may potentially mediate any putative regulation of wild *Drosophila* fitness by the microbiota. It will be interesting in future work to attempt to identify directly the bacterial functions that modulate host gene expression. A related dataset showed that aspects of variation in the bacterial metagenome corresponded to gene expression in wild flies (Bost, Martinson, et al., 2018), suggesting that bacterial functions can indeed be mapped to host gene expression, but causal connections remain to be proven. The fly microbiota are fermentative (Kim et al., 2018; Newell et al., 2014), producing short-chain fatty acids that are expected to modulate the host epigenome (Haws et al., 2020), which we anticipate may underlie host gene expression patterns. Functional characterisation is key to fully understand how microbiota affect host biology (Heintz-Buschart & Wilmes, 2018), but our results accord with other recent analyses of the fly microbiota, which suggest that taxonomy may be a useful proxy for function (Ankrah et al., 2021), and that bacterial order is a functionally informative level of taxonomy (McMullen et al., 2021). Our current results suggest that genus is an informative level, and since genus is nested within order these two sets of findings appear compatible. *Drosophila* lab research suggests that interaction effects can be just as important for host fitness as individual strains (Gould et al., 2018; Ludington, 2022), arguing that a more complete understanding of the system’s biology will be garnered by studying meta-functions of the microbiota because, for example, complex patterns of metabolic cross-feeding among gut microbiota are predicted from just a few species (Ankrah et al., 2021). Ultimately, to distinguish whether bacterial taxonomy is really a sufficient predictor of host function will probably require datasets including paired diversity measurements of both the microbiome and metagenome or metatranscriptome, followed by confirmatory experimental manipulations. *Drosophila’s* experimental utility makes it a strong candidate for this task.

It is important to emphasise that, so far, we have only established correlation between the fly gut microbiota and transcriptome. We have also done so only in a relatively limited number of samples, giving a proof of principle. More work is required to establish causality. Complete control of the microbiota is a considerable strength of the *D. melanogaster* lab model, which provides scope to test further the predictions of surveys of wild animals, to establish causality definitively. Such validation experiments will be valuable because of the potential for confounding variation in correlative surveys (Bost, Martinson, et al., 2018). For example, variation in environmental variables like temperature or humidity may cause changes in the host gut transcriptome and microbiome in concert, yet without a causal relationship between the transcriptome and gut microbiome. Host genotype also modulates both gene expression and microbiome composition (Tabrett & Horton, 2020), across a wide range of species (Chaston et al., 2016; Early et al., 2017; Fuess et al., 2021; Goodrich, Davenport, Beaumont, et al., 2016; Goodrich, Davenport, Waters, et al., 2016; Goodrich et al., 2014; Khan et al., 2019; Kurilshikov et al., 2021; Rajarajan et al., 2022; Small et al., 2019), and in humans can vary among populations and cultures (Syromyatnikov et al., 2022). Host genotype-by-environment effects may also modulate the microbiome (Hahn et al., 2022), though this possibility is under-researched. An additional possibility, which has received relatively little attention, is how host function affects the microbiota: plastic alterations to the gut transcriptome may modulate microbiota composition and abundance in the absence of gene expression that would be recognised as an immune response. Evidence so far suggests that host processes can influence bacterial metabolism (Newell et al., 2022), and this information cannot be detected from 16S and metagenomic surveys. To gain a systematic understanding of these processes will require extensive surveys of wild animals, coupled to careful lab experimentation that manipulates both host and microbe.

While previous lab studies have mostly not addressed relationships between the wild microbiome and transcriptome, laboratory studies do, however, have great utility in addressing a key question that has thus far not been addressed in wild systems: why should microbiota modulate the host transcriptome at all? Two primary functions of gut microbiota are to promote host immunity, and to support host metabolism. Immune signalling pathways are well-established, conserved mechanisms in which extracellular stimuli, such as peptidoglycan and lipopolysaccharide, are detected by receptors which induce signalling events that result in the nuclear import of transcription factors (Lemaitre & Hoffmann, 2007). Immune signalling thus provides a canonical means for the microbiota to modulate host gene expression, but it is not the only possible mechanism. A more cryptic influence may be exerted by modulating the host epigenome (Krautkramer et al., 2016, 2017), altering the standing activity of the host transcriptional machinery, and its responsiveness to stimuli. Lab studies in mammals suggest that a range of epigenetic mechanisms are at play, because no study so far has been able to explain all observed changes in gene expression by one modality of epigenomic regulation. Krautkramer et al (Krautkramer et al., 2016) showed that post-translational modifications of histone proteins are altered in the absence of the microbiota, in a range of mouse tissues. This is expected to alter chromatin accessibility (Buenrostro et al., 2013; Haws et al., 2020), yet Camp et al (Camp et al., 2014) were unable to discern microbiota-dependent changes in chromatin accessibility that corresponded to changes of gene expression in mouse guts. Pan et al (Pan et al., 2018) found genes that were differentially-expressed in the mouse gut, but not all were marked by corresponding changes to DNA methylation. Axenic flies also show signatures of altered gene regulation (Dobson et al., 2016), but any causal epigenetic regulation cannot be due to altered DNA methylation, since flies do not methylate DNA. Altogether this incomplete picture suggests that more scrutiny is required to identify causal mechanisms systematically. One critical factor may be that distinct gut tissues show distinct transcriptomic responses to the microbiota (Sommer et al., 2015), and so finer-scaled analyses of tissue-specific and cell-type-specific responses may be illuminating. Single-cell and multi-omic analyses likely have roles to play, coupled to detailed knowledge of the microbiota themselves. *D. melanogaster* is a strong candidate for providing this fundamental information, given the capacity to couple investigation of fundamental epigenetic mechanisms in laboratory systems with an understanding of naturally-occurring variation in the wild, and the conservation of mechanisms of cell type-specification between flies and other organisms (Dutta et al., 2015). The correspondence between gut microbiota taxonomy and gut transcriptome in wild *D. melanogaster* constitutes an important step along this path.

## Acknowledgments

This work was supported by a UKRI Future Leaders Fellowship (MR/S033939/1) and a University of Glasgow Lord Kelvin Adam Smith Fellowship to AJD. We thank David Sannino, Diana Marcu, Rita Ibrahim and Miriam Wood for helpful discussion, Daniel Haydon for comments on the manuscript, and David McGuinness and John Cole for bioinformatic support. We thank Angela Douglas for supporting the project, assistance with accessing data, and comments on the manuscript.

## Supplementary materials

**Supplementary text.** Description of analytical differences between (Bost, Franzenburg, et al., 2018) and the present study.

**Figure S1. Detailed view of sample collection analysis pipeline.** Figure integrates steps taken by between (Bost, Franzenburg, et al., 2018) and the present study.

**Figure S2. Principal components analysis of OTUs reveals orthogonal major axes of variation in wild fly microbiota. (A)** Biplot showing rotation of OTUs on first two principal components (PCs).

**(B)** Variance explained by each PC. **(C)** Lack of correlation between PCs indicates orthogonal axes of microbiome variation. Pairs plot showing positions of each sample on pairs of PCs, for all pairwise combinatitons. Panels show positions on PCs shown to left (Y axis) and below (X axis), e.g. top-right panel shows positions on PC1 and PC5.

**Figure S3. Mapping microbiome PC values to axes of variation in OTU occurence and abundance**. Major axes of variation in OTU counts were identified by principal components analysis. Resulting PCs were mined for correlations with starting OTU table, identifying OTUs putatively underlying each PC. Panels **A-D** show heatmap analysis for OTUs associated to PCs 1-4, respectively. Heatmaps show values for each PC (line plots at top) and OTU counts (natural log). Sample identities are given at the bottom. Samples are ordered by microbiota PC values (at top). Rows are ordered by hierarchical clustering (dendrograms to right of figures).

**Figure S4. Biological process GO terms in the host gut transcriptome associated with taxonomic variation in the wild fly microbiota.** Gene ontology enrichment was calculated for the full set of all genes associated to microbiome PCs 1-4. Bubble plots show enrichment of GO terms per PC and per ontology. X axis shows P-values (Fisher’s test) for enrichment. GO terms are given on Y axis, ordered by X axis values. Bubble size shows number of genes in the set associated with the GO term, colour shows ratio of n. observed/expected genes.

**Figure S5. Molecular function GO terms in the host gut transcriptome associated with taxonomic variation in the wild fly microbiota.** Gene ontology enrichment was calculated for the full set of all genes associated to microbiome PCs 1-4. Bubble plots show enrichment of GO terms per PC and per ontology. X axis shows P-values (Fisher’s test) for enrichment. GO terms are given on Y axis, ordered by X axis values. Bubble size shows number of genes in the set associated with the GO term, colour shows ratio of n. observed/expected genes.

**Figure S6. Cellular component GO terms in the host gut transcriptome associated with taxonomic variation in the wild fly microbiota.** Gene ontology enrichment was calculated for the full set of all genes associated to microbiome PCs 1-4. Bubble plots show enrichment of GO terms per PC and per ontology. X axis shows P-values (Fisher’s test) for enrichment. GO terms are given on Y axis, ordered by X axis values. Bubble size shows number of genes in the set associated with the GO term, colour shows ratio of n. observed/expected genes.

**Figure S7. Distinct host functions associated to orthogonal axes of microbiota variation**. Gene ontology enrichment was calculated for gene sets associated to microbiome PCs 1-4, for biological process (BP), cellular component (CC) and molecular function (MF) ontologies. Panel shows enrichment per gene set (columns) and ontology (rows). Bubble plots show enrichment of GO terms per PC and per ontology. X axis shows P-values (Fisher’s test, −log_10_ scale) for enrichment. GO terms are given on Y axis, ordered by average of X axis values across all gene sets. Bubble size shows number of genes in the set associated with the GO term, colour shows ratio of n. observed/expected genes. All GO terms per each gene set are shown. See Supplementary Tables 7-10 for detail of GO terms, and Figure 5 for an equivalent plot of the top 5 GO terms per gene set.

**Supplementary table 1.** Library statistics: Dada2 quality control stats from microbial 16S rRNA sequence processing, and RNAseq library properties before and after read filtering

**Supplementary table 2.** DESeq2 analysis of genes associated to microbiota PC1

**Supplementary table 3.** DESeq2 analysis of genes associated to microbiota PC2

**Supplementary table 4.** DESeq2 analysis of genes associated to microbiota PC3

**Supplementary table 5.** DESeq2 analysis of genes associated to microbiota PC4

**Supplementary table 6.** GO enrichment analysis: union of sets of differentially expressed genes associated to microbiota PCs 1-4

**Supplementary table 7.** GO enrichment analysis: differentially expressed genes associated to microbiota PC 1

**Supplementary table 8.** GO enrichment analysis: differentially expressed genes associated to microbiota PC 2

**Supplementary table 9.** GO enrichment analysis: differentially expressed genes associated to microbiota PC 3

**Supplementary table 10.** GO enrichment analysis: differentially expressed genes associated to microbiota PC 4

## Data accessibility and benefit sharing

**Data accessibility statement** Raw data are available from NCBI SRA, with accession numbers PRJNA381755 (16S amplicon sequences) and PRJNA393828 (transcriptomes). Data were originally deposited by Bost et al., (2018).

**Benefits generated** Benefits from this research stem from the sharing of our results and international discussion among colleagues.

