## Supplementary figures and images for "Orthogonal axes of microbiome variation associated with functionally distinct transcriptomic signatures in the gut of wild *Drosophila melanogaster*"

### supplementary figure 1

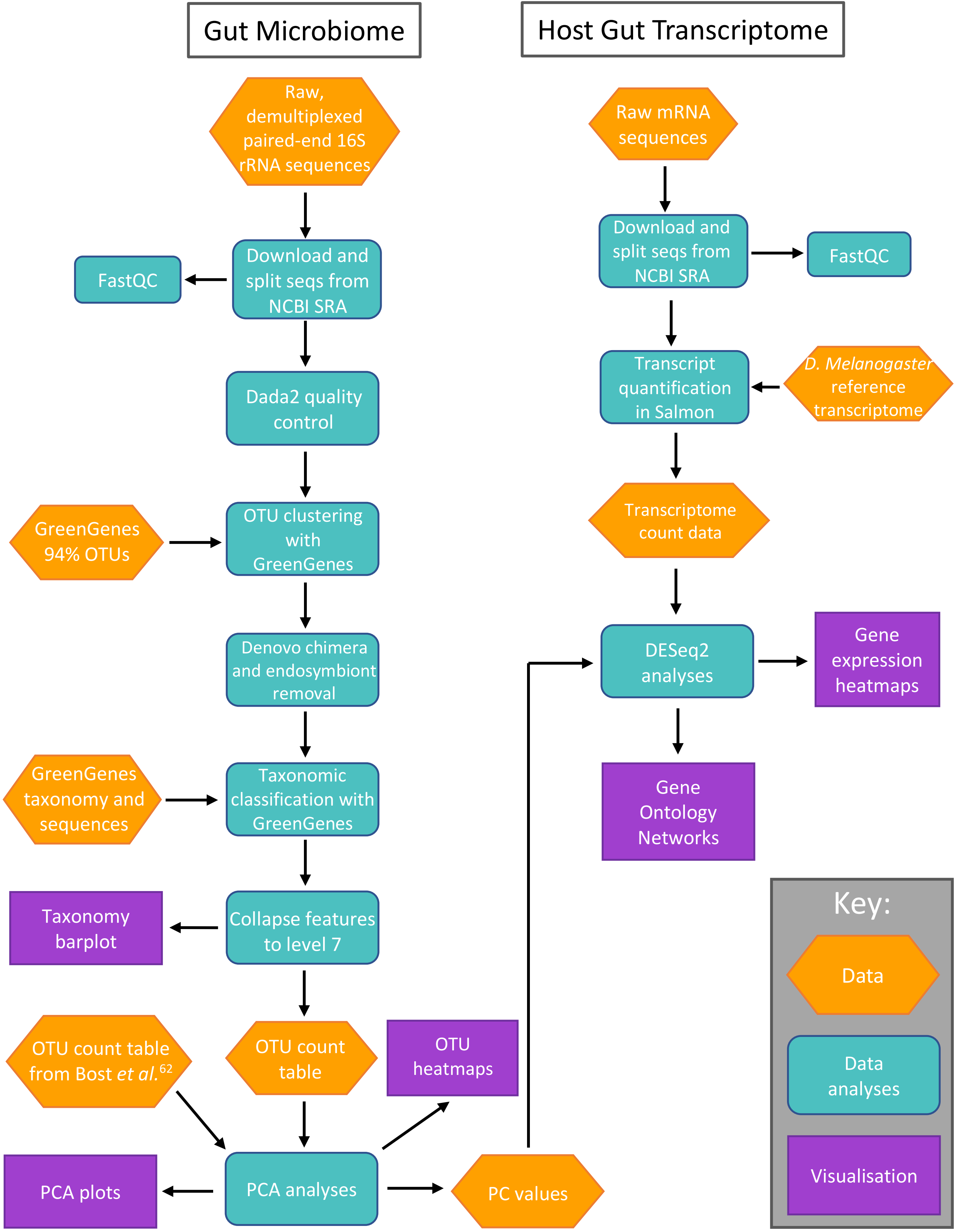

### supplementary figure 2

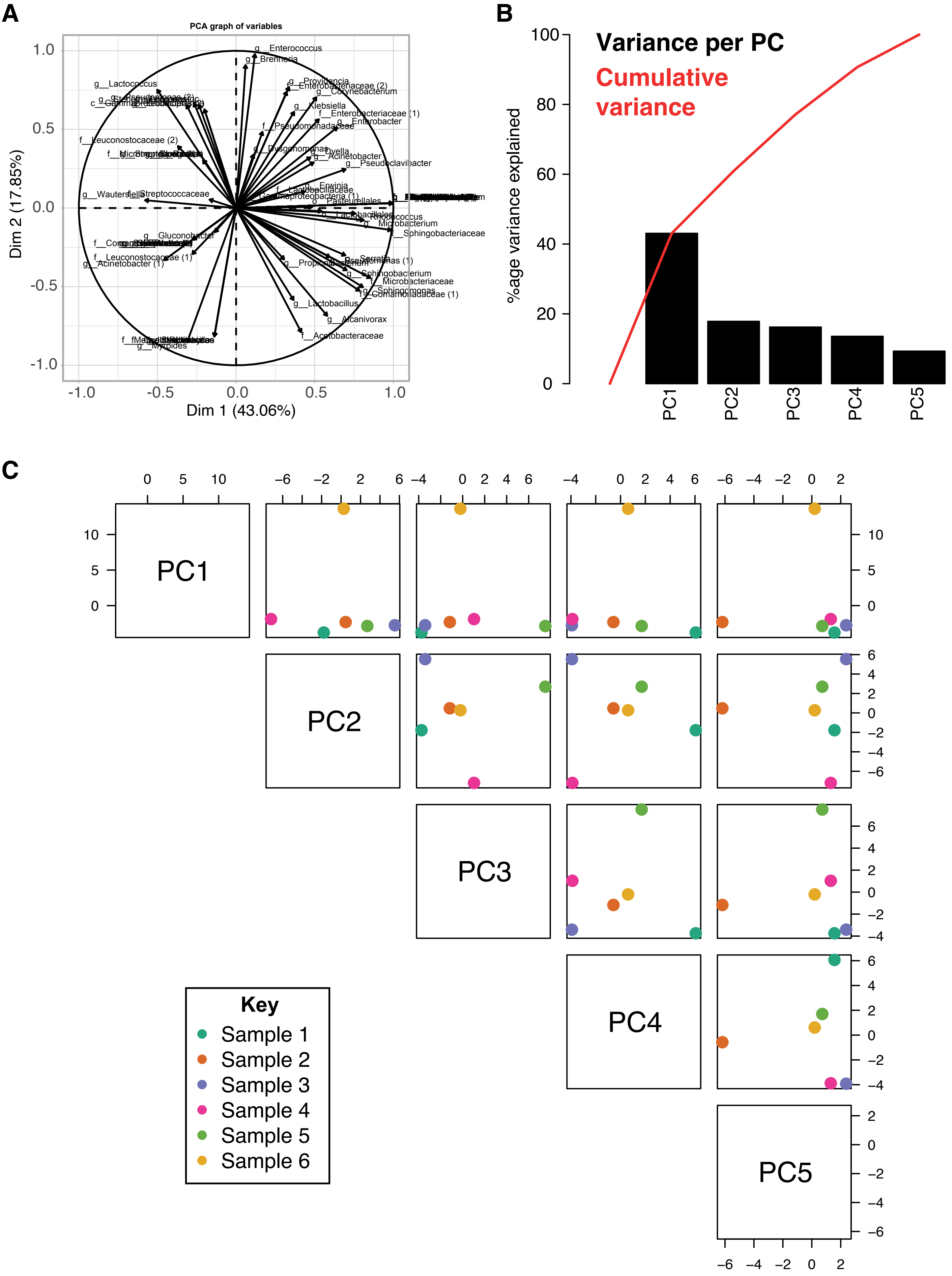

### supplementary figure 3

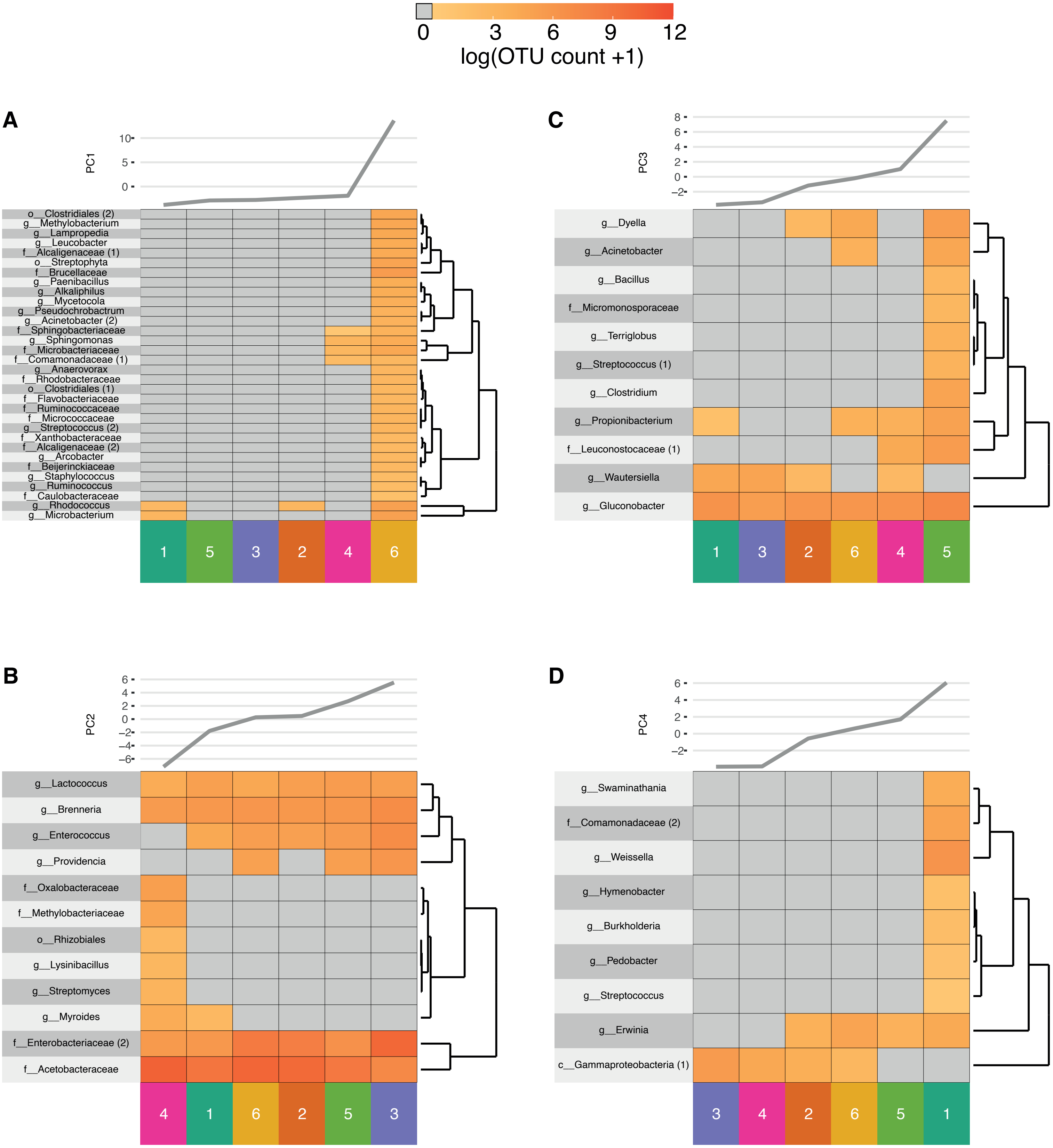

### supplementary figure 4

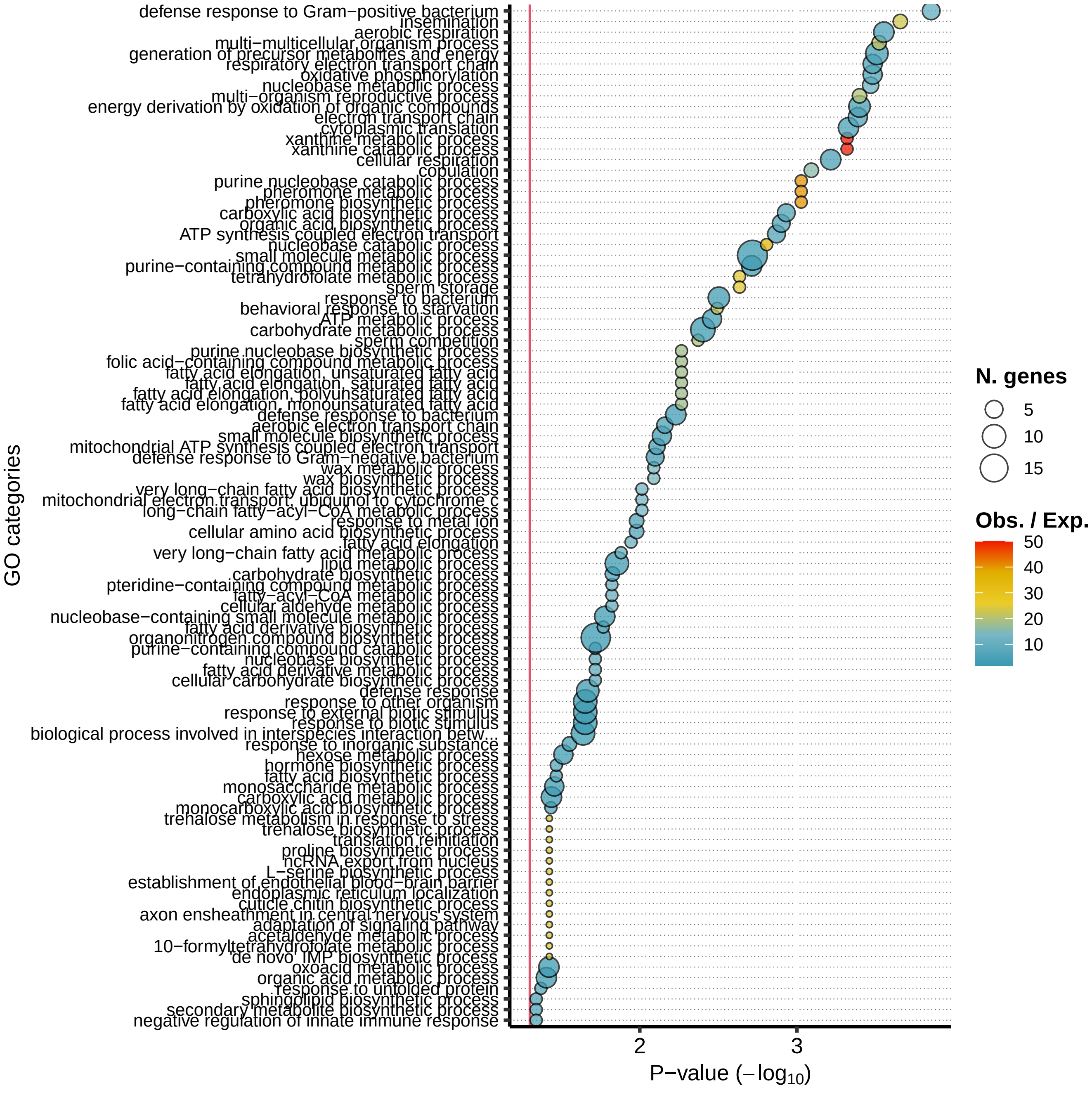

### supplementary figure 5

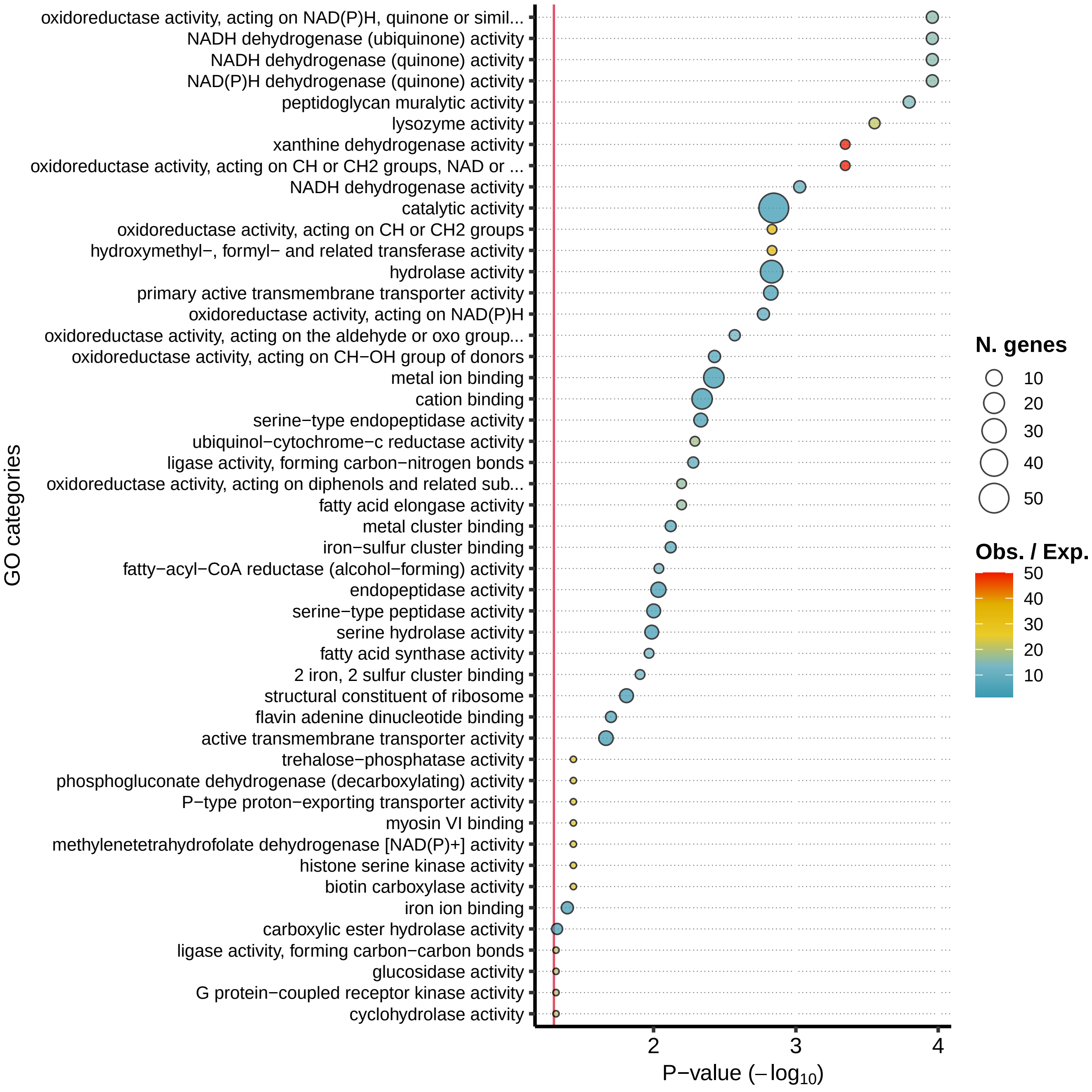

### supplementary figure 6

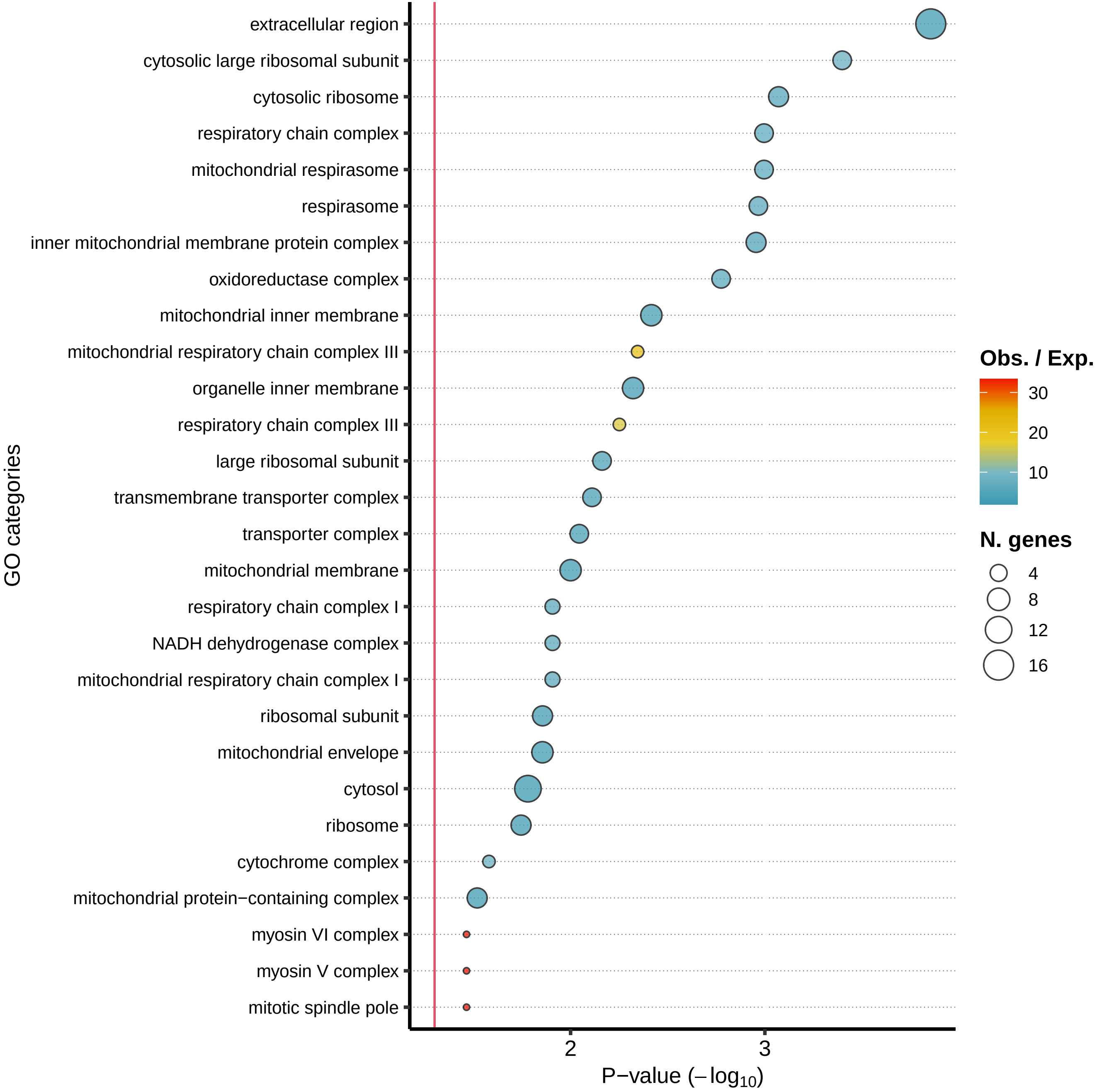

### supplementary figure 7

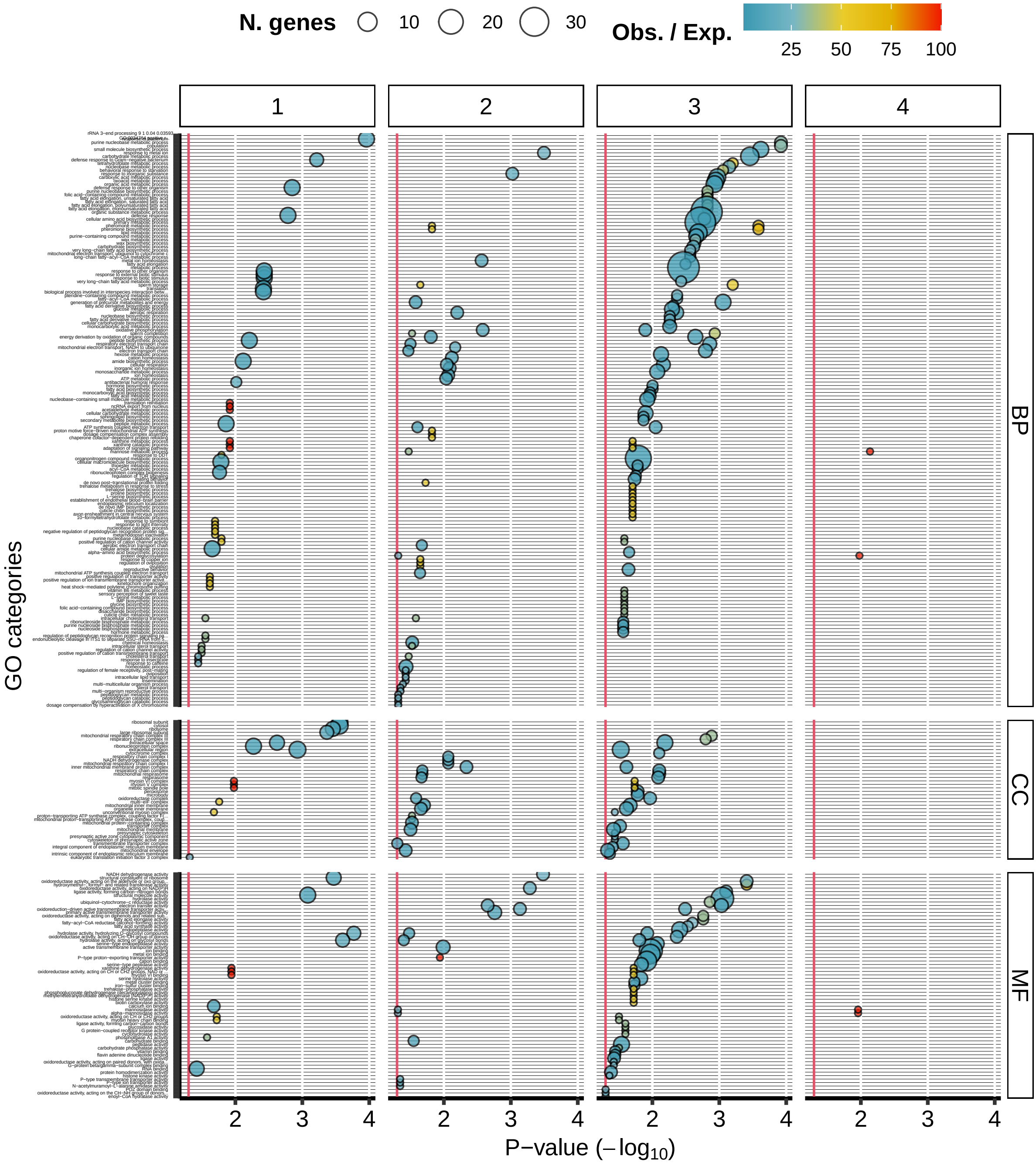
