## supplementary text for "Orthogonal axes of microbiome variation associated with functionally distinct transcriptomic signatures in the gut of wild *Drosophila melanogaster*"

Previously, (Bost et al., 2018) sampled wild male *Drosophila melanogaster* guts, and investigated the relationship between taxonomic composition of the microbiota and the host gut transcriptome. While both the microbiota and transcriptome varied in these samples, little overall covariation detected. This muted correspondence may have been due to a number of factors. This may be due to biological or technical factors. Biologically, diverse microbes may be functionally redundant with respect to the host gut transcriptome, in which case taxonomic variation in the gut microbiota could be irrelevant to the gut transcriptome. Technically, a more sensitive computational analysis may be required. The preceding analysis studied gut transcriptome diversity with a limited set of ~6,414 genes. However the *D. melanogaster* genome contains >17,000 genes, and so detection sensitivity may have been limited. Furthermore, RNAseq data have a number of features (Love et al., 2014) which have motivated the development of specialised tools: applying these tools may enhance sensitivity. Instead, microbiome-transcriptome associations were tested using procrustes analysis and correlation analysis. Procrustes analysis will detect only correspondence between major axes of variation. The sample collections varied by season and geographic site, which may generate transcriptome variation that could dominate impacts of microbial variation, but this does not preclude the potential impact of microbial variation. When correlation analysis was applied, an onerous penalty for multiple hypothesis testing should have been incurred, by testing for correlations between each host transcript and each bacterial operational taxonomic unit (OTUs). The resulting statistical corrections may have increased false negative rate. In addition, the correlation analysis tested for rank correlation: however with the non-normal distribution of count data, and a limited number of samples (6), rank correlations may not reveal the full complexity of the association.

Bost, A., Franzenburg, S., Adair, K. L., Martinson, V. G., Loeb, G., & Douglas, A. E. (2018). How gut transcriptional function of Drosophila melanogaster varies with the presence and composition of the gut microbiota. *Molecular Ecology*, *27*(8), 1848–1859. <https://doi.org/10.1111/mec.14413>

Love, M. I., Huber, W., & Anders, S. (2014). Moderated estimation of fold change and dispersion for RNA-seq data with DESeq2. *Genome Biology*, *15*(12), 550.
